# Identification of suitable reference genes for qPCR expression analysis on the gonads of the invasive mussel *Limnoperna fortunei*

**DOI:** 10.1101/835447

**Authors:** Juliana Alves Americo, Luana Tatiana Albuquerque Guerreiro, Tayssa Santos Gondim, Yasmin Rodrigues da Cunha, Inês Julia Ribas Wajsenzon, Luana Ferreira Afonso, Giordano Bruno Soares-Souza, Mauro de Freitas Rebelo

**Author notes:** Corresponding author: (MFR).

## Abstract

*Limnoperna fortunei* — popularly known as the Golden mussel — is an aggressive invasive species that has been causing environmental damage and adversely affecting economic sectors dependent on freshwater ecosystems in South America. As a non-model species, knowledge about its biology is very limited, especially molecular mechanisms that contribute to its invasiveness, such as its high reproduction rate. Quantitative PCR (qPCR) is considered the gold standard technique to determine gene expression levels and its accuracy relies on the use of stably expressed reference genes for data normalization and to minimize technical variability. Our goal was to identify reliable reference genes to perform gene expression analysis on the gonads of *L. fortunei*. The stability of five candidate genes (*RPS3, EF1a, HS6ST3B, NAPA* and *UBE2F*) in the gonads of male and female mussels was evaluated by using two algorithms, Bestkeeper and Genorm. Results show that *NAPA*, *UBE2F* and *RPS3* genes are stable enough to compose a reliable normalization factor for gene expression analyses comparing both sexes. *HS6ST3B* and *NAPA* were found to be more stable in female gonads; thus, their application as a normalization factor is preferable for studies limited to female processes only.

## 1. INTRODUCTION

Invasive species are estimated to be responsible for at least 39% of the extinctions that have occurred over the past four centuries and are considered the second most important threat to worldwide biodiversity after biological resource use (overexploitation) [1,2]. The golden mussel (*Limnoperna fortunei*), a species native to Asia, has become an aggressive invader in South America, disrupting freshwater ecosystems and causing major economic losses. This mussel (Bivalvia, Mytilidae) causes such extensive changes in the environment — by decreasing substrate availability for native species and by altering the composition and recycling of nutrients in water — that it is considered an ecosystem engineer of freshwater environments, altering their structure and function [3]. Golden mussel infestation affects several economic sectors, including the harvesting and treatment of freshwater for drinking water, fishing and aquaculture, shipping, and especially hydroelectric power generation. So-called “biofouling” of hydroelectric power plant components, such as intake grids, pipes, and pumps, requires that power generation be interrupted for maintenance more frequently. *L. fortunei* is reported to impact at least 40% of Brazil’s hydroelectric power plants (HPPs), causing losses on the order of USD 120 million per year, just in lost revenue [4].

To date, the available control methods, mostly physical (e.g. mechanical removal) or chemical (e.g. chlorine), have been unsuccessful in controlling the invasion by this organism. The primary evidence of the ineffectiveness of these methods is that the golden mussel continues to disseminate on the continent. Since it was first reported in South America in 1991 in the estuary of Argentina’s La Plata River, *L. fortunei* has spread to four other countries: Uruguay, Paraguay, Bolivia, and Brazil. In the Brazilian Pantanal wetlands, the golden mussel is close to the Téles Pires River, one of the affluents of the Amazon basin, threatening the aquatic biodiversity of this vast ecosystem [5]. After almost 30 years of invasion and migration, *L. fortunei* has **spread 5000 km northward**, reaching the Brazilian Northeast [6].

In order to develop more effective methods for controlling the golden mussel, gaps in our knowledge about its biology, especially the mechanisms underlying the biological attributes that make it such a successful invader, need to be narrowed. Certainly key among these biological advantages is its rapid reproduction, which is, in part, a consequence of its rapid development, whereby *L. fortunei* reaches sexual maturity in just three or four months [7], [8]. The golden mussel is dioecious, i.e. sexes are separate, although there are rare reports of hermaphroditism. Reproduction, which is external, occurs almost continuously throughout the year. Gametes are released into the water, where fertilization takes place [7,8].

An important milestone for understanding the biology of this species was the sequencing of its genome [9]. In addition, transcriptomic (RNA-seq) analyses have been performed **for tissues such as the digestive glands, foot, mantle, muscle [9, 10, 11], and**, more recently, for tissue from the gonads of both sexes [12]. This latest work characterized the gene expression profile of adult male and female gonads and revealed almost 4000 sex-biased transcripts in this tissue. Despite the value of these genome-wide data resources, experimental assessment of gene functions in the golden mussel is incipient.

RNA-seq studies produce informative large-scale gene expression data without the need for any previous sequence knowledge. Nevertheless, quantitative PCR (qPCR) **remains the gold standard technique for accurate determination of gene expression [13,14]. Indeed**, as the RNA-seq cost per sample is still prohibitive for large sample sizes, **qPCR is favored to validate** specific RNA-seq expression results [15], as well as to investigate the functions of specific genes in broader experimental conditions.

The accuracy of qPCR assays is largely dependent on the use of reference genes (RGs) stably expressed in these experimental conditions. RG expression values are used to normalize the data obtained from genes of interest, in order to adjust for technical variation in the qPCR workflow as, for example, in the quality and quantity of input samples and in the retrotranscription efficiency [14,16]. Therefore, suitable reference genes that have had their stability validated under the study conditions, are essential to identify changes in gene expression which are truely of biological origin.

Nevertheless, the use of suitable reference genes has not been the rule in the literature. Recently, a review of gene expression studies in several bivalves of ecotoxicological interest was performed and it was observed that validated RGs were used in less than 40% of the studies. Of these, less than 25% documented the validation process [17]. Another frequent methodological fault is the use of only a single reference gene in 86.3% of the studies, when the use of normalization factors, consisting of two or more validated RGs, has been considered standard practice by the scientific community for over 15 years [14,16].

In this study, we sought to validate RGs for gene expression studies in golden mussel gonads, in order to be able to then investigate the expression pattern of putative genes thought to be involved in the reproduction of this mollusk. RGs considered candidates for such evaluation were selected from available RNA-seq data and from the scarce literature describing the validation of RGs for the gonads of close species. We were able to validate suitable RGs for the normalization of qPCR data produced from male and female golden mussel gonads.

## 2. MATERIALS AND METHODS

### 2.1. Selection of candidate reference genes

Candidate reference genes were selected using transcriptome and transcripts expression data previously generated from the gonads of spawning male and females golden mussels [12]. The expression matrix was used for EdgeR and DESeq2 methods to perform statistical tests between the groups. Transcripts were evaluated according to their fold change (|logFC|), when comparing males and females, and the coefficient of variation (CV%= standard deviation/mean*100) of their expression counts (trimmed mean of M-values, TMM) [18] across the biological replicates (three for each sex). The selection was based on the premise that the more stable the gene, the lower the LogFC and CV% values. Transcripts for which the TMM count was zero in at least one of the samples, as well as those which did not match any gene in the reference genome, [9] were not considered.

In order to avoid polymorphic positions during the primer design process, the identification of SNPs based on the aforementioned RNA-seq dataset was undertaken using a strategy similar to the one described as mpileup-transcriptome [19], except that the transcriptome assembly was generated by a concatenation of three different strategies as described previously [12]. Briefly, the concatenated transcriptome was used as the reference and reads were mapped using Bowtie2 [20]. Then, Samtools mpileup and bcftools [20,21] were used to call SNPs from the mapped reads and a final multifasta file with IUPAC ambiguity codes was generated. These sequences were used for primer design with the Primer-BLAST tool [22], avoiding regions containing SNPs. Primers were designed to span exon-exon junctions, except for *HS6ST3B*, which is a single-exon gene. This was based on the exon-intron structure predicted for each gene in the golden mussel reference genome [9]. The presence of secondary structures (hairpin and dimers) in the primers was evaluated using the Oligoanalyzer tool (IDT). Finally, primers specificity was verified by BLASTn searches against the *L. fortunei* genome (GCA_003130415.1). All primers generated in this study are described in Table 1. All transcript sequences were retrieved from the transcriptome assembly previously performed [12], which are available in the NCBI sequence read archive (SRA) under accession number PRJNA587212. Transcript identities were confirmed by BLASTx (https://blast.ncbi.nlm.nih.gov/Blast.cgi) searches against the UniprotKB/Swiss-Prot database. Amplicon sequences are provided in S1 Table.

**Table 1:**
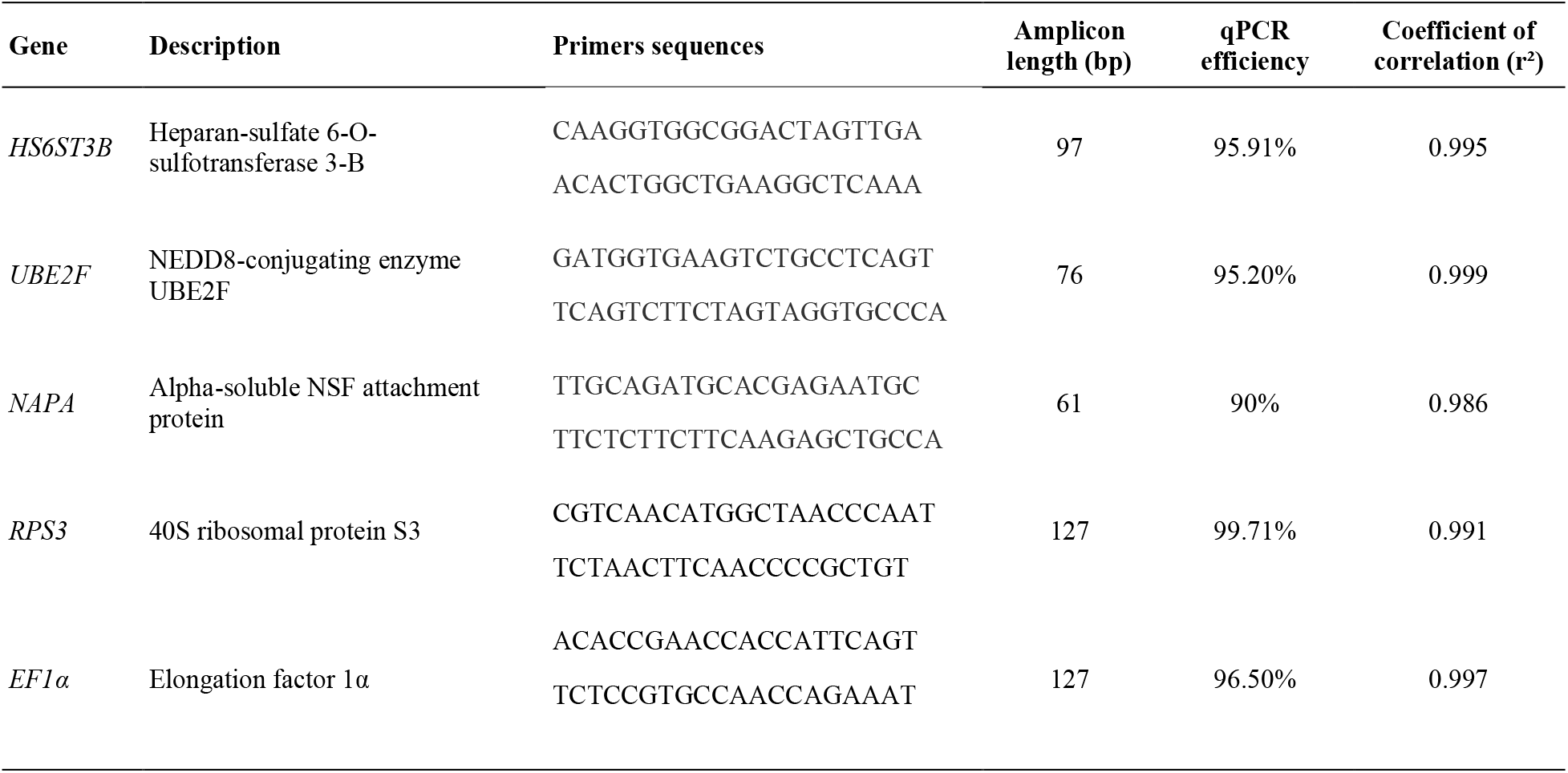
Primers used for qPCR amplification of candidate reference genes evaluated in this study.

### 2.2. Collection, sample processing and histological sex identification

A total of 15 golden mussel (*L. fortunei*) specimens were collected from the Paranapena River in the state of São Paulo, Brazil). *L. fortunei* is an exotic species and is not characterized as endangered or protected species. The gonads of each specimen were dissected and split in two. One half was stored in RNAlater solution (Sigma Aldrich) at 4°C during 24h, after which these specimens were stored at −20°C until further processing. The other gonad half was fixed in paraformaldehyde 4% (Isofar) and stored at 4°C. Fixed gonads underwent standard histological processing steps: dehydration by increasing concentrations of ethanol (Isofar), clarification with xylene (Isofar) and impregnation in Paraplast Plus^®^ (Sigma Aldrich). Histological sections (7 μm) were obtained, stained with hematoxylin and eosin and examined under a light microscope for sex identification and gonadal developmental stage classification according to Callil, Krinski, and Silva [22].

### 2.3. RNA extraction and cDNA synthesis

Next, specimens underwent DNAse digestion using components of the TURBO DNA-free kit (Invitrogen) for the removal of any trace genomic DNA. Then, the absence of DNA contamination and the RNA concentration in the specimens were determined fluorometrically with Qubit™ dsDNA BR Assay and Qubit™ RNA BR Assay Kits (Invitrogen), respectively. The synthesis of cDNA was performed using 600 ng of total RNA of each sample. The retrotranscription reactions were carried out using the High capacity cDNA synthesis kit (Applied Biosystems), following the manufacturer’s protocol.

### 2.4. Quantitative PCR and gene stability analysis

All the qPCR analyses were performed in technical triplicates in 96-well plates on a QuantStudio 3 Real-Time PCR system (Applied Biosystems). The amplification efficiencies of each primer pair were determined by running standard curves with five points of a serial 2-fold dilution (ranging from 10 ng to 0.6 ng of total RNA input per reaction). The PowerUp SYBR Green Master Mix (Applied Biosystems) kit was used to perform the qPCR assays, for which the following components were added in each reaction: 5μl of 2X master-mix, 1 μl of diluted cDNA (10 ng/μl, except for standard curves as detailed above), primers at a final concentration of 0.3 μM (for *UBE2F*, *EF1a* and *RPS3* genes) or 0.6 μM (for *HS6ST3B* and *NAPA* genes) and water up to 10 μl. The reactions were subjected to temperature cycling as follows: 50°C (2 min), 95°C (2 min), then to 40 repeats of: 95°C (15 s), 56°C (15 s), 72°C (1 min). At the end of the amplification, reactions underwent a melting curve analysis and amplicons were analyzed by agarose electrophoresis (2%) with ethidium bromide in order to certify the assay specificity. Negative controls, where no template was added, were performed for all the genes and every 96-well plate. The raw Cq data (consisting of the means of the three technical replicates for each specimen) were used to generate a Box-plot graph using the BoxPlotR tool [23] (available at: http://shiny.chemgrid.org/boxplotr/). Finally, the stability of the candidate reference genes was evaluated using two algorithms: BestKeeper V1 [24] and Genorm [16], with the latter embedded in the Qbase+ software, version 3.2 (Biogazelle, Zwijnaarde, Belgium - www.qbaseplus.com).

## 3. RESULTS AND DISCUSSION

We were able to identify stably expressed RGs in the golden mussel gonads because we started from an available RNA-seq dataset and used qPCR to validate our findings. In order to select the most stably expressed genes in this dataset, we first ordered the transcripts according to the fold change (|logFC|), when comparing males and females, narrowing down to the first 300 transcripts displaying the smallest fold change values. Then, the remaining transcripts were placed in ascending order of CV (%) of the expression counts (TMM) across all the specimens (three for each sex). As summarized in Table 2, three candidate reference genes were selected among the transcripts displaying lower fold change (0.018 - 0.030) and a lower expression Coefficient of Variation (4.83% - 6.76%). As determined by BLAST (Table 2), these transcripts are (1) heparan-sulfate 6-O-sulfotransferase 3-B (*HS6ST3B*), (2) NEDD8-conjugating enzyme UBE2F (*UBE2F*) and (3) alpha-soluble NSF attachment protein (*NAPA*).

**Table 2:** Candidate reference genes BLAST identification and stability measures in the gonad RNA-seq data. The information presented is for the best hit found for each transcript as a result of BLASTx searches performed against the UniProt / Swiss-Prot database. Stability measures, Fold change (Mean |logFC|) and Expression CV% (based on TMM counts) were retrieved from available *L. fortunei* gonad transcriptomes data [12].

| Transcript ID | Best hit Acc N° and Organism | Best hit gene name | Best hit description | E-value | Identity (%) | RNA-seq Fold change | RNA-seq Expression CV% |
| --- | --- | --- | --- | --- | --- | --- | --- |
| GGt_755971_c0_g1_i1 | A0MGZ7.1 (*Danio rerio*) | HS6ST3B | Heparan-sulfate 6-O-sulfotransferase 3-B | 2.00E-100 | 56.99% | 0.03 | 4.83% |
| DNt_685619_c0_g1_i1 | Q969M7 (*Homo sapiens*) | UBE2F | NEDD8-conjugating enzyme UBE2F | 3.00E-77 | 61.41% | 0.02 | 5.83% |
| DNt_712450_c0_g1_i1 | Q9DB05.1 (*Mus musculus*) | NAPA | Alpha-soluble NSF attachment protein | 3.00E-123 | 67.95% | 0.02 | 6.76% |
| DNt_730425_c0_g1_i1 | E2RH47.1 (*Canis lupus familiaris*) | RPS3 | 40S ribosomal protein S3 | 2.00E-155 | 91.45% | 0.28 | 20.39% |
| DNt_467182_c0_g1_i1 | Q9YIC0.1 (*Oryzias latipes*) | EF1α | Elongation factor 1α | 0.00E+00 | 88.80% | 0.11 | 49.33% |

*HS6ST3B* is involved in the heparan sulfate (HS) proteoglycan biosynthetic process, and is part of a gene family composed of three other 6-O-Sulfotransferases found in zebrafish (*Danio rerio*) [25]. HS plays important roles in cell regulation; HS itself is modulated by modifications such as the addition of sulfate groups by this family of genes. In invertebrates such as *Caenorhabditis elegans*, a single gene is usually identified and shown to be essential for normal development [26]. The other two genes are part of distinct processes: *NAPA* is a component of the fusion machinery required for vesicular trafficking. [27,28], while *UBE2F* promotes post-translational protein modification by conjugating the ubiquitin-like protein NEDD8 to its target proteins [29].

Two additional genes were selected based on previous studies conducted on the gonads of other bivalve molluscs: 40S ribosomal protein S3 (*RPS3*) and elongation factor 1α (*EF1α*), which are both involved in protein synthesis. Even though these two genes presented lower stability in the analyzed RNA-seq data as indicated by fold change and TMM CV% values (Table 2), they are among the most frequently used RGs in other bivalves [17]. *RPS3* proved to be stable in the gonads of the mussel *Mytilus galloprovinciallis* [30], while *EF1α* was previously validated as a RG for the gonads of the scallop *Pecten maximus L*. [31], the oyster *Pinctada margaritifera* [32], and the scallop *Mizuhopecten yessoensis* [30,33] and for this reason these genes were selected for evaluation in the golden mussel gonads.

The qPCR assay conditions established for the measurement of these five candidate RGs proved to be efficient, with amplification efficiencies of all genes falling within the recommended range (90-110%) [14], as shown in Table 1. The melting curves and the amplicon migration patterns seen with agarose electrophoresis were consistent with the specific amplification of each target as documented in the supporting material (S1 and S2 Figures), showing that the qPCR assays were also specific to the genes of interest. Thus, after initial qPCR standardization and validation, candidate reference genes were measured in a set of *L. fortunei* gonad specimens, including eight female and seven male specimens. For each sex, there were specimens from three gonad developmental stages as follows: in maturation (1 male, 2 females), mature (1 male, 2 females) and spawning (5 males, 4 females).

For a preliminary assessment of the stability of the genes, Cq data distribution was displayed in a box plot (Fig 1). We first analyzed all the results for the whole dataset (females + males) and then for each sex separately. Each of the three analyses showed a median Cq between 15 and 0, within the recommended range of medium expression level for reference genes [34]. Considering the complete data set, the gene with the greatest difference between the minimum and maximum Cq values was *HS6ST3B* (7.4), while the gene with the smallest difference was *NAPA* (2.7). These results suggest that these two candidates may be the most unstable and stable genes, respectively, when considering specimens of both sexes. Indeed, *HS6ST3B* is the gene for which the box plot data distribution profile for female and male specimens differed the most (Fig 1).

**Fig 1.**
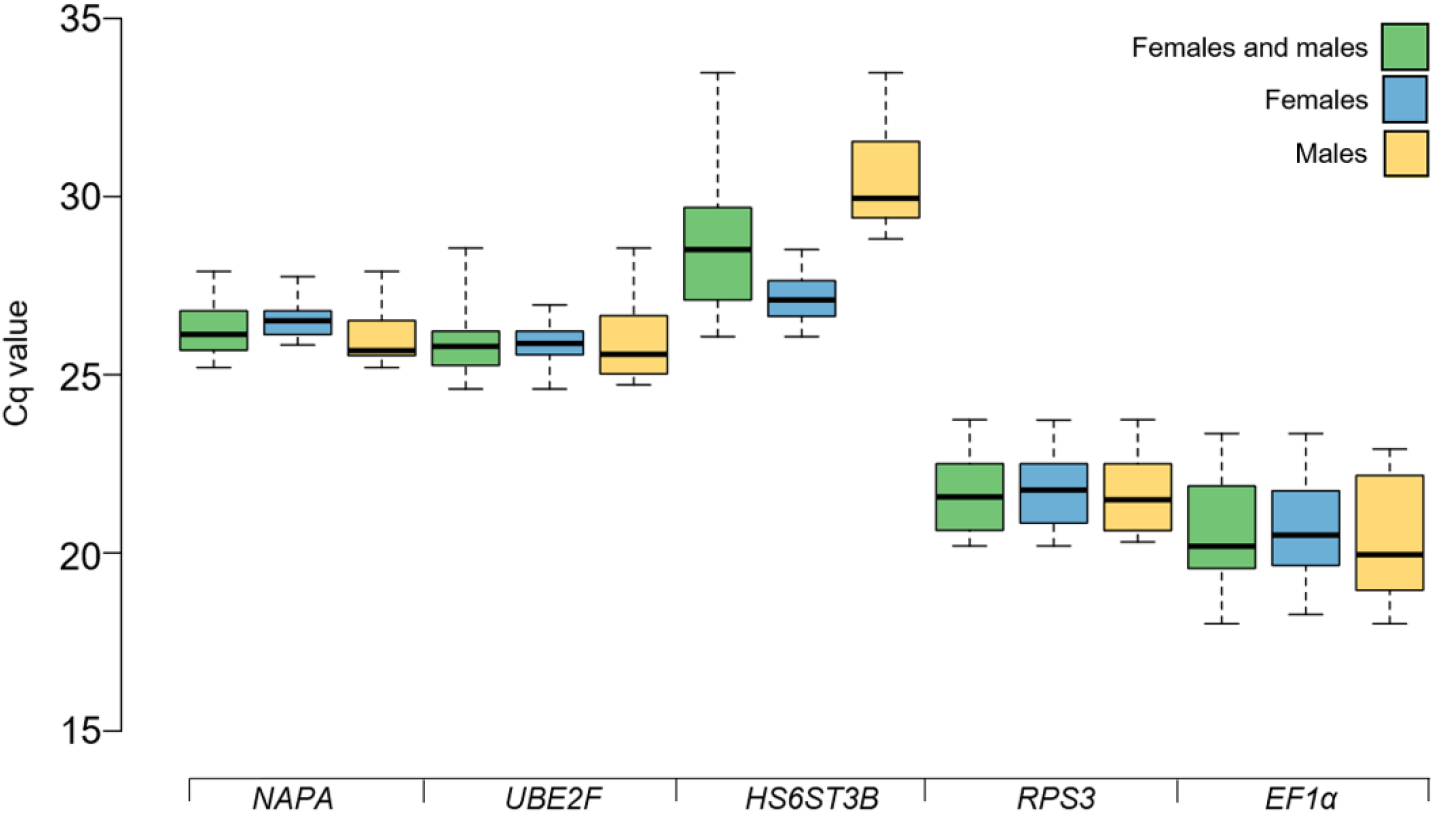
Candidate reference genes expression variation according to Cq values distribution. Center lines show the medians; box limits indicate the 25th and 75th percentiles; whiskers extend to minimum and maximum values.

The stability of candidate RGs should not be assessed solely on the variability of Cq values and, therefore, amplification data were also analyzed using two of the most widely used algorithms to evaluate RGs stability: Genorm [16] and BestKeeper [24]. Genorm is based on the principle that the expression ratio between two perfect reference genes should be identical in all the analyzed samples. In order to evaluate that, this algorithm performs pairwise variation analysis, where the standard deviation (SD) of the expression ratios observed across the samples is calculated for each gene pair combination. A measure of stability (Genorm M value) is calculated for each gene based on the arithmetic average of the pairwise variation values observed for the combination of the gene with each one of the other candidates under analysis. Therefore, the lower the M value, the lower the variability and the higher the gene stability. To rank the genes according to their stability, a stepwise exclusion process was performed, where the least stable gene is removed and the average M value is recalculated for the remaining genes until a single gene is left. The result of this process is shown in Fig 2A, where data for both sexes were analyzed together and separately.

**Fig 2.**
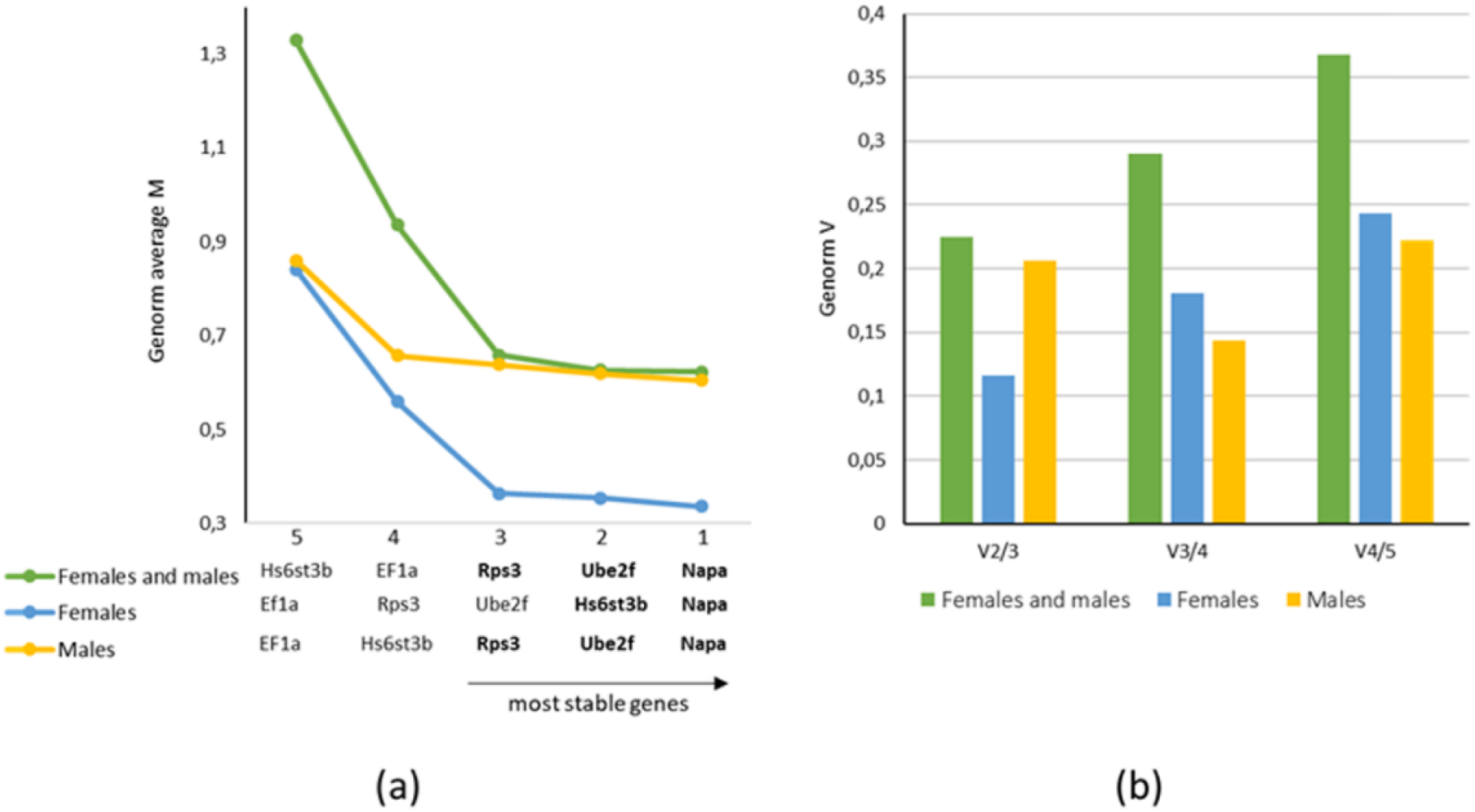
Genorm stability analysis of the candidate reference genes. In (a), the curve shows the stepwise exclusion of the least stable genes and the resulting gene stability rankings for each analyzed scenario. (b) shows the results from pairwise analysis performed to determine the minimal recommended number of reference genes to be applied for data normalization.

The stepwise exclusion process identified the same three genes (*NAPA* > *UBE2F* > *RPS3*) as the most stably expressed in males and in the combined (female and male) set of specimens, while the same process when applied to the female specimens data alone, generated the three top-ranked stable genes (with the lowest average M values) that were different (*NAPA* > *HS6ST3B* > *UBE2F*). In each of the three scenarios, *NAPA* was depicted as the most stable gene, while *EF1a* was consistently among the two least stable genes. Even though *HS6ST3B* was highly stable in females, this gene was among the the two least stable in males and in the combined female + male data, which is consistent with the box plot profiles observed in Fig 1. For homogeneous samples, Genorm M values ranging up to 0.5 are considered ideal, while for heterogeneous and more divergent samples, this cutoff value is less strict and M values up to 1.0 are considered acceptable [35]. This can be considered the case for the gonads of both sexes, a tissue where gene expression is highly influenced by the sex of the organism [36][12]. Indeed, in each of the three analyzed scenarios, the three top-ranked genes all showed average M values below the 1.0 threshold.

In order to evaluate the minimal number of Reference Genes to be used for accurate gene expression, normalization factors (RGs Cq geometric mean) for increasing numbers of genes were calculated, and the pairwise variation (V) between two sequential combinations were computed (Vn/n+1). In this analysis, a larger variation means that the addition of another gene could be beneficial. However, once V is below 0.15, an additional gene is not required [16]. Fig 2B shows that for females and males, separately, the use of two (*NAPA* and *HS6ST3B*) and three (*NAPA*, *UBE2F* and *RPS3*) genes respectively, is sufficient to reach V < 0.15. However, for the complete dataset, a V value lower than this threshold was not reached, and the lowest observed V value was 0.22 (V2/3). The 0.15 threshold should not be considered too strictly, and for difficult samples the normalization by the three most stable genes is generally recommended [16].

The second algorithm used to evaluate gene stability was BestKeeper [24], which like Genorm, is based on the principle that stably expressed genes should retain some correlation between their expression levels across the samples representative of the conditions of interest. In order to evaluate that, BestKeeper computes a series of descriptive statistics (Table 3A) and performs a Pearson’s pairwise correlation analysis (Table 3B) of each gene with each other and between each gene and a Bestkeeper Index (BKI). The BKI is calculated as the geometric mean of the candidate RGs Cq values in each specimen. The coefficient of correlation (r), the coefficient of determination (r^2^) and the p-value permit us to evaluate how well every gene correlates with the BKI, a mean measure of expression levels of the analyzed candidate RGs.

**Table 3:** BestKeeper results for the five evaluated candidate Reference Genes (RG). In section A, the descriptive statistics are shown. In section B, the repeated pairwise correlation analysis with the five candidate RGs are shown, while in section C, the results of the same analysis with only the three most stable genes in each scenario are shown. F=Female; M=Male.

| Gene | NAPA |  |  | UBE2F |  |  | RPS3 |  |  | HS6ST3B |  |  | EF1a |  |  |
| --- | --- | --- | --- | --- | --- | --- | --- | --- | --- | --- | --- | --- | --- | --- | --- |
| Sex | F + M | F | M | F + M | F | M | F + M | F | M | F + M | F | M | F + M | F | M |
| *A: Descriptive statistics* |  |  |  |  |  |  |  |  |  |  |  |  |  |  |  |
| SD [± CP] | 0.68 | 0.45 | 0.85 | 0.77 | 0.47 | 1.15 | 0.98 | 0.87 | 1.09 | 1.70 | 0.58 | 1.44 | 1.46 | 1.24 | 1.69 |
| CV [% CP] | 2.59 | 1.68 | 3.27 | 2.97 | 1.83 | 4.41 | 4.51 | 4.00 | 5.01 | 5.92 | 2.15 | 4.72 | 20.5 | 1.24 | 8.27 |
| geo Mean [CP] | 26.34 | 26.55 | 26.11 | 25.92 | 25.85 | 26 | 21.69 | 21.74 | 21.65 | 28.69 | 27.16 | 30.55 | 20.57 | 20.62 | 20.36 |
| ar Mean [CP] | 26.36 | 26.56 | 26.13 | 25.94 | 25.86 | 26.03 | 21.73 | 21.77 | 21.68 | 28.76 | 27.17 | 30.59 | 18.01 | 20.68 | 20.45 |
| min [CP] | 25.2 | 25.84 | 25.2 | 24.6 | 24.6 | 24.72 | 20.19 | 20.19 | 20.31 | 26.07 | 26.07 | 28.81 | 23.35 | 18.27 | 18.01 |
| max [CP] | 27.9 | 27.75 | 27.9 | 28.55 | 26.96 | 28.55 | 23.74 | 23.73 | 23.74 | 33.47 | 28.51 | 33.47 | 7.08 | 23.35 | 22.91 |
| *B: Repeated pairwise correlation analysis (N=5 genes)* |  |  |  |  |  |  |  |  |  |  |  |  |  |  |  |
| coeff. of corr. [r] | 0.828 | 0.877 | 0.957 | 0.880 | 0.809 | 0.917 | 0.921 | 0.927 | 0.983 | 0.717 | 0.941 | 0.975 | 0.856 | 0.854 | 0.923 |
| coeff. of det. [r<sup>2</sup>] | 0.686 | 0.769 | 0.916 | 0.774 | 0.654 | 0.841 | 0.848 | 0.859 | 0.966 | 0.514 | 0.885 | 0.951 | 0.733 | 0.729 | 0.852 |
| P value | 0.001 | 0.004 | 0.001 | 0.001 | 0.015 | 0.004 | 0.001 | 0.001 | 0.001 | 0.003 | 0.001 | 0.001 | 0.001 | 0.007 | 0.003 |
| *C: Repeated pairwise correlation analysis (N=3 genes)* |  |  |  |  |  |  |  |  |  |  |  |  |  |  |  |
| coeff. of corr. [r] | 0.942 | 0.964 | 0.977 | 0.934 | 0.938 | 0.967 | 0.956 | - | 0.964 | - | 0.964 | - | - | - | - |
| coeff. of det. [r<sup>2</sup>] | 0.887 | 0.929 | 0.955 | 0.872 | 0.880 | 0.935 | 0.914 | - | 0.929 | - | 0.929 | - | - | - | - |
| P value | 0.001 | 0.001 | 0.001 | 0.001 | 0.001 | 0.001 | 0.001 | - | 0.001 | - | 0.001 | - | - | - | - |

First, genes were ranked according to the standard deviation (SD) of Cq values (the lower, the better), which is considered the most relevant BestKeeper statistic. Genes presenting high SD Cq values are excluded, with the acceptable cutoff in the literature ranging from 1.0 [24] to 1.5 [37]. In the gene rankings for the group combining female and male specimens (*NAPA* > *UBE2F* > *RPS3* > *EF1a* > *HS6ST3B*) and the group containing only male specimens (*NAPA* > *RPS3* > *UBE2F* > *HS6ST3B* > *EF1a*) the two least stable genes, with the highest SD Cq values (Table 3A) were *EF1a* and *HS6ST3B*, while the group consisting of female specimens (*NAPA* > *UBE2F* > *HS6ST3B* > *RPS3* > *EF1a*) the two least stable genes were *RPS3* and *EF1a*.

Next, the two least stable genes in each scenario were excluded, and pairwise correlation analysis were repeated, in order to access the correlation between each remaining gene and the BKI, which was recalculated based on the three most stable genes only (Table 3C). Finally, genes were ranked according to the coefficient of correlation (r) observed in this final pairwise correlation analysis. The same three most stable genes previously found by Genorm for the female + male dataset (*RPS3* > *NAPA* > *UBE2F*), as well as for females (*NAPA*=*HS6ST3B* > *UBE2F*) and males (*NAPA* > *UBE2F* > *RPS3*) were confirmed by BestKeeper, differing only in the rank order of the genes. Additionally, another difference was that *NAPA* and *HS6ST3B* were found in this last analysis to be equally stable in the female gonad specimens.

Therefore, Genorm and BestKeeper results are in agreement, reinforcing that among the investigated candidates RGs, *NAPA*, *RPS3* and *UBE2F* are the three most stable genes for the gonads of the golden mussel, and should be applied together to establish a reliable normalization factor, regardless of the sex of the mussel. On the other hand, the results also show that *HS6ST3B* is particularly stable in females, and together with *NAPA*, seems to be sufficient for accurate qPCR data normalization in studies tracking gene expression in female gonad specimens only.

*RPS3*, one of the RGs selected from the literature, turned out to be among the three most stable genes. However, it proved to be less stable than the RGs picked from RNA-seq data. Although there is no universal reference gene, and validation is always required for the tissues and species of interest, *RPS3* has shown some consistency, being stable in the gonads of several other bivalves, as mentioned earlier, and might be considered as a candidate to be validated for the gonads of other mussel species.

The results of this study should be seen in light of several limitations. The first is that the analyzed specimens were limited to the developmental stages where the sexes have already differentiated and can be observed through the histological analysis of the gonads. Therefore, for gene expression studies in juveniles, when sex is not yet evident, validation of the genes evaluated in the present work, and possibly other candidates, is required. Another limitation is that the stability of the genes evaluated here was verified only in the gonads and, therefore, these reference genes should not be applied to studies that aim to determine the differential expression between this organ and other tissues without additional validations.

To our knowledge, this is the first work that sought to validate RGs for the golden mussel in an effort to implement robust qPCR assays for gene expression analysis in this non-model organism. The application of qPCR to this species has been limited to the detection of larvae in the environment by absolute quantification, for which RGs are not required [38–40] and to one research article where qPCR was applied to evaluate gene expression in this mussel foot, but using a single non-validated RG [11]. We hope that the data presented here may help to change this picture, enabling the execution of reliable gene expression studies in the golden mussel gonads and contributing to advance the knowledge of the molecular mechanisms of key aspects of its extremely efficient reproduction, one of the distinguishing features of this aggressive invader.

## 4. CONCLUSIONS

In this study, we evaluated candidate reference genes for normalization of qPCR data generated from the gonad specimens of male and female golden mussels. Among the investigated genes, *RPS3*, *UBE2F* and *NAPA* were found to be the most stable and are indicated for studies with individuals of both sexes or only males. *HS6ST3B* and *NAPA* genes showed greater stability in females and should be preferentially used to compose a normalization factor in studies investigating only female gonad specimens.

## 5. SUPPORTING INFORMATION

**S1 Fig. Melting curves of the candidate reference genes (*RPS3, EF1a, HS6ST3B, NAPA* and *UBE2F*)**. In each graph, the red lines correspond to the “no template controls” and, therefore, as expected, show no signal of amplification.

**S2 Fig. Ethidium bromide agarose (2%) electrophoresis analysis of PCR products**. Lane 1: 50bp Ladder (Sinapse Inc, M1041); Lane 2: *NAPA*; Lane 3: *UBE2F*; Lane 4: *HS6ST3B*; Lane 5: *RPS3*; Lane 6: *EF1a*; Lane 7: 100bp Ladder, Ready-To-Use (Sinapse Inc, M106).

**S1 Table. Transcript region amplified by qPCR (amplicon sequence) using the primers described in this study**.

