## Supplementary material for "Identification of suitable reference genes for qPCR expression analysis on the gonads of the invasive mussel *Limnoperna fortunei*": S1 Fig

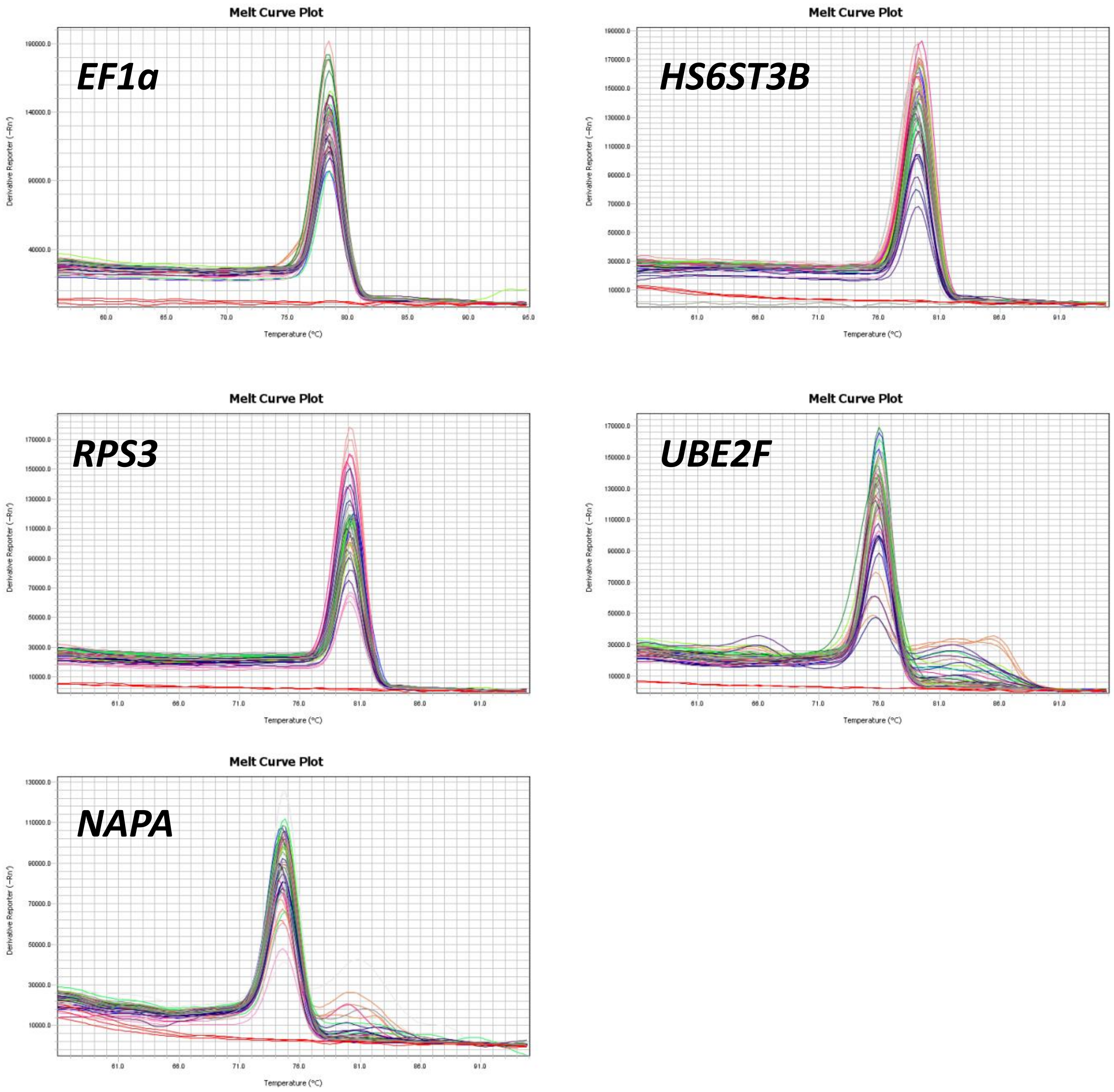


**S1 Figure: Melting curves of the candidate reference genes (RPS3, EF1α, HS6ST3B, NAPA and UBE2F)**. In each graph, the red lines correspond to the "no template controls" and, therefore, as expected, show no signal of amplification.
