## Supplementary material for "Identification of suitable reference genes for qPCR expression analysis on the gonads of the invasive mussel *Limnoperna fortunei*": S2 Fig

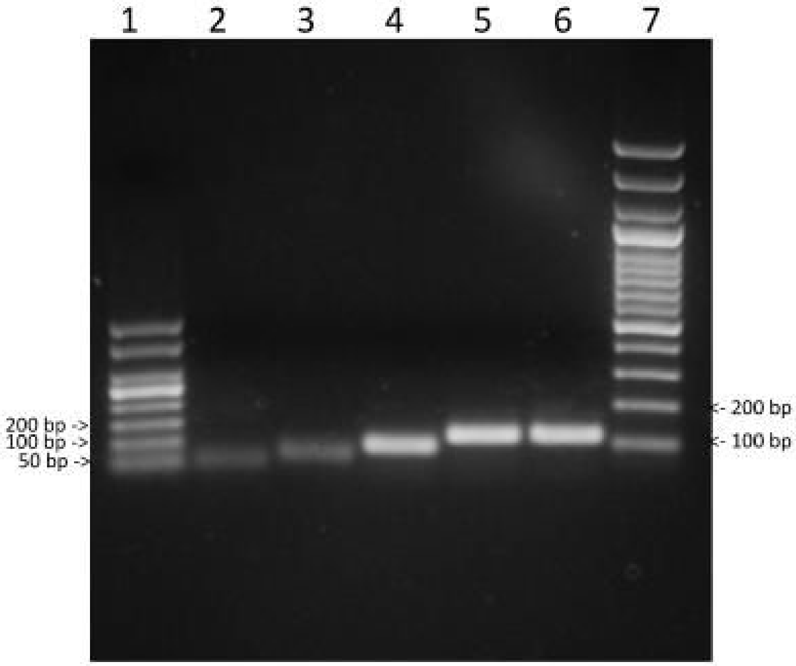


**S2 Figure: Ethidium bromide agarose (2%) electrophoresis analysis of PCR products.** Lane 1: 50bp Ladder (Sinapse Inc, M1041); Lane 2: *NAPA*; Lane 3: *UBE2F*; Lane 4: *HS6ST3B*; Lane 5: *RPS3*; Lane 6: *EF1a*; Lane 7: 100bp Ladder, Ready-To-Use (Sinapse Inc, M106).
